# Top-down attention modulates auditory sustained responses but not the neural processing advantage for vowels

**DOI:** 10.64898/2026.06.10.731417

**Authors:** E.V. Orekhova, K.A. Fadeev, M.V. Morozova, I.V. Romero Reyes, A.M. Plieva, T.A. Stroganova

## Abstract

Temporal and spectral regularities characteristic of vowels are preferentially encoded by the auditory system, giving rise to a prolonged enhancement of a negative electromagnetic response in the auditory cortex - sustained negativity (SN). However, it remains unclear how sustained attention to the auditory stream interacts with this intrinsically enhanced processing. Here, we used MEG to examine SN and its differential response to vowel-like acoustic patterns versus spectrally complex noise, while modulating subject’s attention to the auditory stimulation.

Four stimulus types (600 ms duration; vowel-like sounds with and without pitch, periodic non-vowel sounds, and aperiodic noise) were presented to 30 adults during passive listening and an active gap-detection task. In 16% of randomly selected trials, a 50-ms silent gap was inserted into the stimuli, which served as a target. The number of non-target trials between successive targets ranged from 1 to 10, with the highest probability corresponding to an interval of six trials. Only non-target trials were analyzed.

Analysis of reaction times and omission errors indicated that target expectancy increased with the number of trials since the previous target. Task-related attention enhanced an early orienting-related negativity (∼50–150 ms) across all stimulus types, irrespective of target expectancy. It also accelerated the processing of periodicity in non-vocal sounds but did not affect the processing of vowels. In contrast, the later segment of the SN (150–600 ms) was strongly modulated by proximity to the previous target, decreasing immediately after target events and increasing as the likelihood of target occurrence rose. Despite these attentional and expectancy-related modulations, the enhanced neural processing of periodic vowels relative to noise remained stable.

These findings indicate that sustained attention modulates both early and late SN responses and accelerates the extraction of periodicity in non-vocal sounds, yet does not alter the neural advantage conferred by vowel-related regularities.

## 1 Introduction

The auditory cortex is capable of remarkably rapid extraction of sensory information from conspecific vocal sounds. The ultra-early emergence of vocal sound–specific responses is consistent with selective tuning of cortical neurons, which exhibit early-onset and sustained neuronal activity to restricted combinations of acoustic features (Walker, Bizley, King, & Schnupp, 2011; Wang, Lu, Snider, & Liang, 2005), effectively acting as a filter for biologically salient acoustic patterns (Nelken, 2008). Within this framework, response properties are not fixed but may instead be shaped by experience and attentional state (Atiani, Elhilali, David, Fritz, & Shamma, 2009; Schneider, et al., 2021).

In the human auditory cortex, responses to a prolonged sound (hundreds of milliseconds) are manifested as a series of transient components superimposed on a sustained negative shift of electric current termed sustained negativity (SN) (Gutschalk, Patterson, Scherg, Uppenkamp, & Rupp, 2004; Orekhova, et al., 2024). Converging evidence demonstrates that SN is extremely sensitive to auditory patterns (“auditory figures”), which enhance the negative shift relative to acoustically matched noise (Barascud, Pearce, Griffiths, Friston, & Chait, 2016; Sohoglu & Chait, 2016; Southwell, et al., 2017), giving rise to sustained negativity associated with processing of acoustic regularities, i.e. sustained processing negativity (SPN) (Fadeev, et al., 2024). Notably, vowel sounds elicit especially robust SPN (Fadeev, et al., 2024; Gutschalk & Uppenkamp, 2011; Hewson-Stoate, Schönwiesner, & Krumbholz, 2006; Orekhova, et al., 2024).

This vowel-specific SPN emerges as early as ∼50 ms after stimulus onset (Edmonds, et al., 2010; Hewson-Stoate, Schönwiesner, & Krumbholz, 2006), and persists throughout stimulus presentation. The earliest part of this vowel-related increase in negativity has been linked to rapid extraction of acoustic features in lower-level auditory cortex, such as formant structure and periodicity (Edmonds, et al., 2010; Mizuochi, et al., 2005; Yrttiaho, Tiitinen, Alku, Miettinen, & May, 2010), whereas its later phase may reflect higher-order phonetic processing, including vowel categorization (Bidelman, Moreno, & Alain, 2013) and subsequent monitoring of a stable auditory object (Gutschalk & Uppenkamp, 2011). Although SN enhancement (i.e. SPN) to vowels is robust during passive listening, suggesting a high degree of automaticity, the extent to which it is modulated by attention remains unclear.

Because SN has a high signal-to-noise ratio and is similarly modulated by vocal sounds in both adults and children (Orekhova, et al., 2024), it provides a promising measure for probing different stages of auditory speech processing in clinical populations, including auditory processing disorders, autism spectrum disorder (ASD), and dyslexia. Critically, however, the interpretation of SN differences in such populations depends on understanding the mechanisms underlying its enhancement.

Although SN advantage for vowels reflects the intrinsic salience of these stimuli, an important unresolved question is whether and to what extend this advantage depends on enhanced attention allocated to the auditory stream in which they occur. This question is particularly important in clinical contexts, where atypical attentional engagement is a common feature of many neurodevelopmental and psychiatric disorders and may influence auditory processing from early stages. As compared to non-speech sounds, speech sounds are known to engage greater sustained attention to the auditory stream in typically developing individuals (Jaramillo, et al., 2001; Zhang, Li, Chen, & Gong, 2018), but less consistently in clinical populations (Ceponiene, et al., 2003; Whitehouse & Bishop, 2008). Importantly, attentional engagement may operate across multiple processing stages—from early sensory gain modulation to later attentional guidance of perceptual analysis.

Previous studies of auditory SN have demonstrated that attention modulates responses to temporally regular auditory patterns: directing selective attention toward specific stimulus features enhances an early SN (Chait, de Cheveigné, Poeppel, & Simon, 2010; de la Chapelle, et al., 2025; Mittag, et al., 2013), whereas diverting attention away from the auditory stream attenuates SN from approximately 50 ms onwards (Molloy, Lavie, & Chait, 2019; O’Sullivan, Shamma, & Lalor, 2015). Taken together, these findings suggest that the enhanced processing of auditory patterns is possible without attention being directed to the sound, but that the perception of the auditory object is facilitated by attention.

However, these effects have been established primarily for novel, artificial acoustic regularities. In contrast, vowel sounds are highly familiar and behaviorally relevant auditory objects whose representations are deeply embedded in cortical networks (Belin, Zatorre, Lafaille, Ahad, & Pike, 2000). It therefore remains unclear whether attention exerts a comparable influence on such well-learned stimuli, or whether their processing advantage is largely stimulus-driven.

In particular, it is unknown whether sustained, non-selective attention to an auditory stream—without explicit demands on phonetic discrimination—modulates vowel processing, and whether such modulation can alter the previously described “vowel-specific” effects. Addressing this issue is critical for distinguishing between genuinely altered phoneme-level processing and differences in sustained attention to vocal streams, which may occur in clinical populations, such as e.g. ASD.

In the present study, we used MEG and source modeling to investigate auditory-cortical SN responses during passive listening and sustained attention. To identify which acoustic features underlying vocal-sound processing are influenced by attention, we presented three types of test stimuli: regular vowels containing both formant structure and periodicity, vowel-like sounds lacking periodicity, and spectrally complex periodic sounds lacking formant structure. Spectrally complex noise stimuli containing neither periodicity nor formant structure served as controls. Under passive condition, subjects watched a silent movie while stimuli (600-ms periodic and aperiodic vowels, periodic non-vowel sounds, and energy-matched noise) were presented though the earphones. In the active listening condition, the same stimuli were presented, but participants were instructed to monitor them for a rare target event—a brief temporal gap that could occur within any of the presented sounds. Because the probability of target occurrence increased with the number of standard trials since the preceding target, target expectancy was assumed to increase systematically across trials, thereby modulating sustained attentional engagement. Importantly, none of the sounds served as targets themselves; rather, each stimulus constituted a monitored foreperiod during which a target gap could occur. To minimize contamination from target-related activity and isolate neural effects associated with sustained attention and target expectancy, only non-target trials were included in the analyses. This design allowed us to investigate how fluctuations in sustained attention over time influence the processing of vowels, their derivatives, and control noise stimuli, and whether the preferential neural encoding of the spectral and temporal regularities characteristic of vowels is affected by attention.

## 2 Methods

### 2.1 Participants

The study included 30 healthy adults (14 women) aged 20.2 to 40.9 years (M = 26.8 ± 5). None of the participants had a known history of neurological or psychiatric disorders.

The study was approved by the Ethics Committee of the Moscow State University of Psychology and Education. All participants provided written informed consent after receiving a complete explanation of the experimental procedures.

### 2.2 Stimuli

Stimuli were synthetic vowel-like sounds developed based on a paradigm previously described in studies by Gutschalk and Uppenkamp (2011) and Uppenkamp et al. (2006). However, the formant structure of our stimuli more closely matched Russian vowel phonemes /a/, /i/, /o/, /e/, and /□/. The vowel phonemes /a/, /i/, /o/, /e/ have close counterparts in English (e.g., /a/ as in *father*, /i/ as in *machine*, /o/ as in *note*, /e/ as in *bed*), whereas /□/ has no direct English equivalent.

Stimuli were constructed from damped sinusoids with an exponentially decaying envelope. This envelope ensured rapid decay of the sinusoidal components and produced a temporal structure resembling vocal fold pulsations during phonation (Irino & Patterson, 1996). To generate regular periodic vowel-like sounds, the damped sinusoids were repeated with a period of 12 ms, corresponding to a fundamental frequency of 83 Hz, which falls within the typical range of the male voice. The total duration of each stimulus was 600 ms, and the sampling rate was 44.1 kHz.

The carrier frequencies of the sinusoidal components in the vowel stimuli corresponded to the four lowest formants selected within the typical adult male range. For the vowel /a/, the formant frequencies were F1 = 730 Hz, F2 = 1090 Hz, F3 = 2400 Hz, and F4 = 3400 Hz; for /i/, F1 = 270 Hz, F2 = 2280 Hz, F3 = 3000 Hz, and F4 = 3360 Hz; for /o/, F1 = 520 Hz, F2 = 625 Hz, F3 = 2450 Hz, and F4 = 3220 Hz; for /□/, F1 = 300 Hz, F2 = 1480 Hz, F3 = 2230 Hz, and F4 = 3270 Hz; and for /e/, F1 = 440 Hz, F2 = 1800 Hz, F3 = 2550 Hz, and F4 = 3450 Hz. These formant configurations provided stable vowel perception, as confirmed in a pilot test involving five naive adult participants. These stimuli are hereafter referred to as “*periodic vowels*.”

In addition to the periodic vowel stimuli, three additional stimulus types were generated by disrupting periodicity, formant structure, or both.

In the “*non-periodic vowel*” condition, periodicity was disrupted by temporally randomizing the repetition of the damped sinusoids: the position of each sinusoid was randomly shifted within a range of ±6 ms relative to its corresponding position in the periodic vowel stimulus. This procedure was performed independently for each of the four formants and resulted in the loss of perceived periodicity and the associated perception of voice pitch, while preserving the overall formant structure of the stimulus. Subjectively, these stimuli were perceived as vowels with a breathy or hoarse voice quality.

In the “*periodic non-vowel*” condition, the carrier frequency of each successive damped sinusoid was randomly selected from a set of eight possible frequencies distributed on a geometric scale within a range of ±0.5 octaves relative to the original formant frequency. Frequency selection was performed independently for each formant, with the additional constraint that two consecutive sinusoids could not have the same carrier frequency. This procedure disrupted the stable formant structure while preserving periodicity, resulting in stimuli that were perceived as non-speech sounds resembling the “musical rain effect previously described in (Uppenkamp, Johnsrude, Norris, Marslen-Wilson, & Patterson, 2006).

In the “*non-periodic non-vowel*” condition, which serves as a control condition, both periodicity and formant structure were disrupted using the procedures described above.

The stimuli, which were characterized by temporal or frequency regularities (vocal pitch or formant structure)—periodic vowels, non-periodic vowels, and periodic non-vowels—were considered test stimuli. The stimulus lacking any regularity (non-periodic non-vowel sounds) was a ‘control’ stimulus.

All stimuli were low-pass filtered with a cutoff frequency of 5000 Hz. This procedure reduced the influence of high-frequency components on the subjective perception of loudness. For all stimuli, the onset rise time was around 0.5 ms. The stimuli used in this study are available as the *Supplementary material*.

Target stimuli consisted of modified versions of the above sounds containing a 50 ms acoustic gap. The gap onset was pseudo-randomly selected within an interval from 150 to 400 ms after stimulus onset. To avoid spectral artifacts and audible clicks, both the onset and offset of the gap were smoothed using 5 ms cosine ramps. The gap stimuli were perceived as a temporary interruption of a continuous sound rather than as a separate acoustic event, and were clearly distinguishable from both the uninterrupted standard stimuli and the interstimulus intervals.

### 2.3 Procedure

Auditory stimuli were presented binaurally through plastic sound-conducting tubes inserted into the ear canals. The tubes were secured to the MEG helmet to prevent possible noise caused by contact with the participant’s clothing. Stimulus intensity was set to 90 dB SPL.

The experiment consisted of two consecutive sessions: (1) a passive listening session (12 min) and (2) the gap detection task (16 min). The order of the sessions was counterbalanced across participants. Figure 1 illustrates the experimental design.

**Figure 1.**
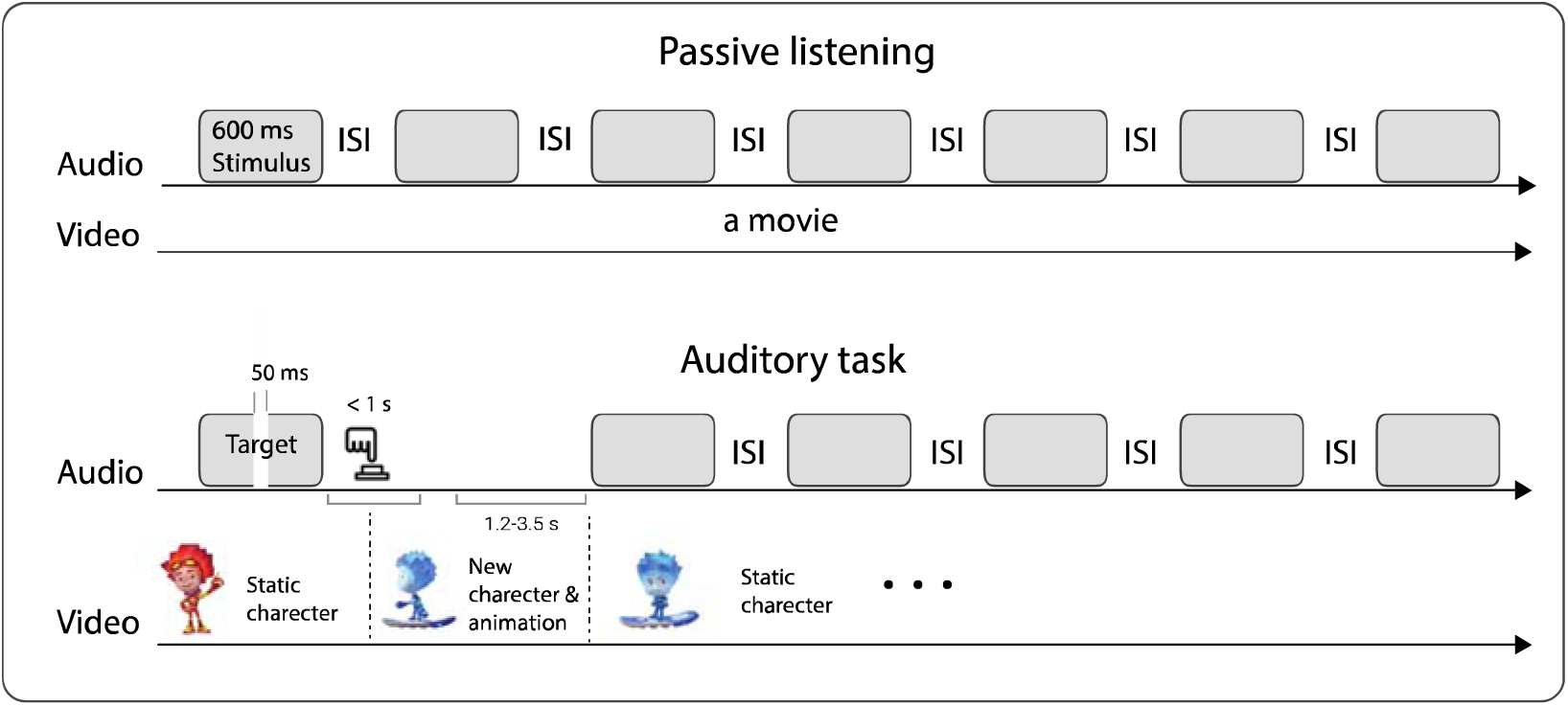
Experimental design. During the *passive listening* condition, four types of 600-ms stimuli were presented in random order, with inter-stimulus intervals (ISIs) varying between 500 and 800 ms, while participants watched a movie of their choice. During the *auditory task*, the same auditory stimuli were presented while a static cartoon character was displayed on the screen. Participants detected the target: a 50-ms gap that could occur within any stimulus type with equal probability (∼14% of all stimuli). A correct response (button press) triggered the appearance of a new character and a short animation, after which the same character remained static until the next correct response.

During the passive listening session, 136 stimuli of each type (approximately 27 for each of 5 vowels or their derivatives), were presented at random with inter-stimulus intervals varied within 500-800 ms. Participants were instructed to watch a silent video (a movie or cartoon of their choice) and to ignore the auditory stimuli.

During the gap detection task, the same stimuli were present, but some of them (N = 90; approximately 14%) contained a 50-ms gap. Participants were instructed to press a button whenever they detected a gap. The target (gap) could occur within any stimulus type with equal probability, ensuring that all stimulus categories were equally task-relevant. The number of standard stimuli of each of the four types was the same as in the passive condition (N = 136) and they occurred at random.

Figures 2A and 2B show the distribution of target-event probability as a function of the number of trials since the previous target, along with the corresponding hazard function representing the instantaneous likelihood of target occurrence. Targets never directly followed the previous target; they were extremely rare during the second trial after the target. Thereafter, the target probability increased, peaking at the seventh trial after the target, and then gradually decreased. The likelihood of target appearance increased nearly exponentially with each target trial after the target.

**Figure 2.**
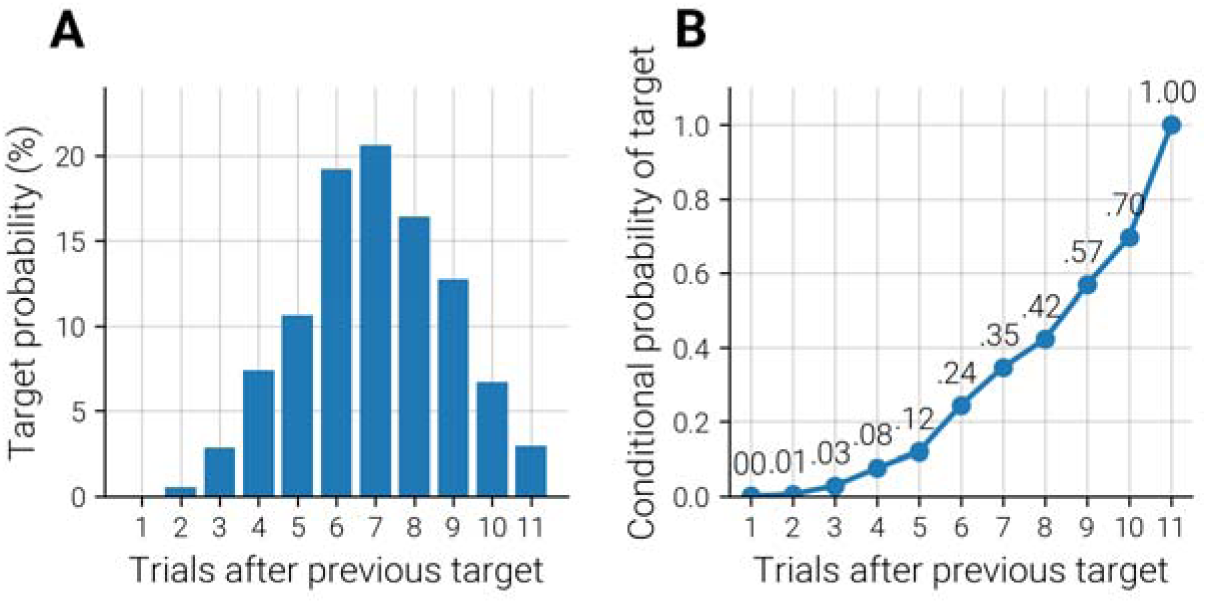
Target-event probability (A) and instantaneous likelihood (B) as a function of trial number following the previous target.

**Figure 3.**
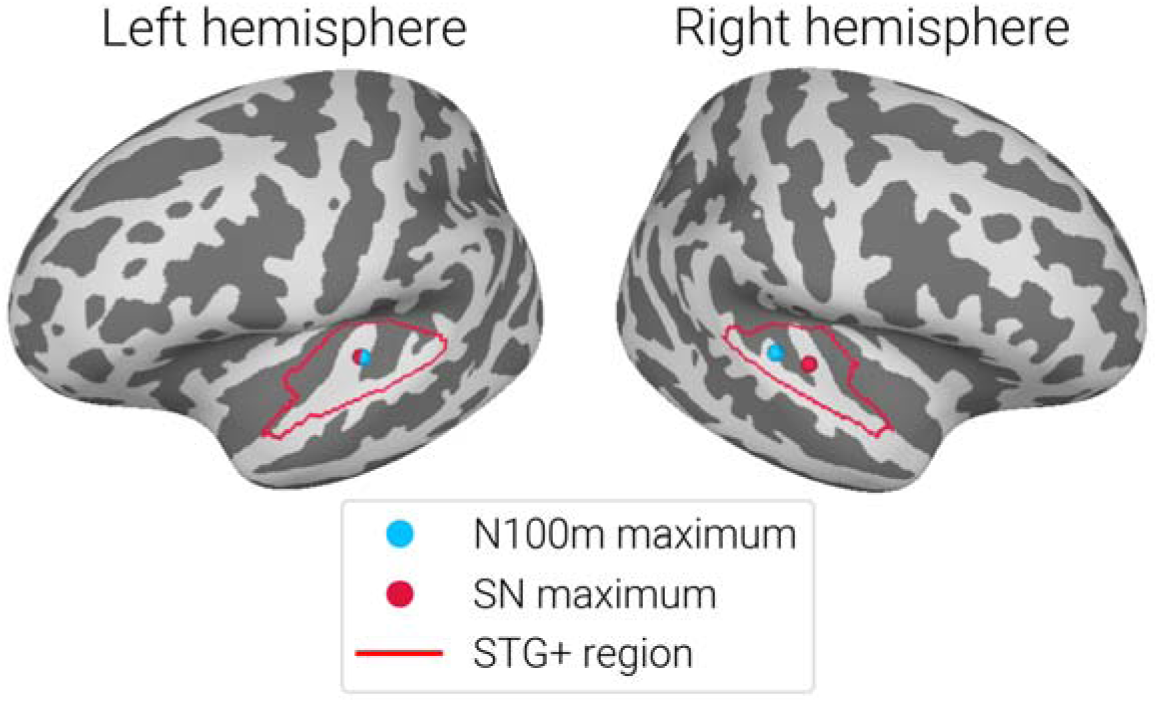
Regions of the auditory cortex (STG+) where individual ‘N100m’ and ‘late SN’ ROIs were defined. Borders of these regions are shown on the FS-average brain for demonstration purposes, but subjects’ ROIs were defined using individual brain models. Blue and red dots indicate, respectively, the average coordinates of the vertices with maximal amplitude of N100m and of SN in the 300–600 ms interval, where the SN component does not overlap with transient evoked activity.

In the gap detection task, a static character from a popular cartoon movie was displayed on the screen during auditory stimulation. To maintain participant engagement, each correct response triggered the new cartoon character. The new character was animated for 700–2700 ms and then remained static until the next correct response. The animations varied in both character identity and animation type, with a total of 84 unique animations used. If the participant failed to respond within 1.4 s after target onset, a large red diagonal cross was displayed on a black background for 1 s followed by a new trial. However, all responses exceeding 1 s were considered misses.

### 2.4 Data acquisition and preprocessing

All participants underwent structural MRI scanning with acquisition of T1-weighted images. Individual cortical reconstructions and cortical parcellations were performed using FreeSurfer version 8.1.0.

MEG data were recorded at the Moscow Center for Neuro-cognitive Research (MEG-Center) using Elekta VectorView Neuromag 306-channel MEG detector array (Helsinki, Finland) with 0.1 - 330 Hz filters and 1000 Hz sampling frequency. Bad channels were visually detected and labeled, after which the signal was preprocessed with MaxFilter software (v.2.2) in order to reduce external noise using the temporal signal-space separation method (tSSS) and to compensate for head movements by repositioning the head in each time point to an “optimal” common position (head origin). This position was chosen individually for each participant as the one that yielded the smallest average shift across all data epochs after motion correction.

Further preprocessing steps were performed using MNE-Python (v1.9) [https://mne.tools?utm_source=chatgpt.com]. The data were filtered using notch filters at 50 and 100 Hz and a low-pass filter at 110 Hz. Periods in which peak-to-peak signal amplitude exceeded thresholds of 7 × 10^-12^ T for magnetometers or 7 × 10^-10^ T/m for gradiometers, as well as “flat” segments in which signal amplitude was below 1 × 10^-15^ T for magnetometers or 1 × 10LJ¹³ T/m for gradiometers, were automatically excluded from further processing.

Signal-space projection (SSP) was applied to correct for cardiac and eye-movement artifacts based on ECG, vertical EOG (vEOG), and horizontal EOG (hEOG) recordings. We subsequently excluded data segments where head rotation exceeded 10°/s along any of the three spatial axes, head velocity exceeded 4 cm/s in 3D space, or head displacement exceeded 20 mm relative to the ‘optimal’ head position.

The cleaned data were then epoched from −0.2 to 0.8 s relative to stimulus onset. The mean number of clean standard epochs per participant and stimulus type was 133 (range: 122–136) and 132 (range: 75–152) for the task and passive conditions, respectively. Epochs were baseline-corrected by subtracting the mean signal amplitude within the −200 to 0 ms prestimulus interval.

### 2.5 MEG data analysis

To obtain the source model, the cortical surfaces reconstructed with FreeSurfer were represented by dense triangulated meshes containing approximately 130,000 vertices per hemisphere. The cortical surfaces were subsequently downsampled to a source grid of 4,098 vertices per hemisphere, corresponding to an average spacing of approximately 4.9 mm between adjacent source points along the cortical surface.

Source reconstruction of the event-related fields was performed using standardized low-resolution brain electromagnetic tomography (sLORETA) (Pascual-Marqui, 2002). The forward solution was computed using a single-layer boundary element model (BEM) based on the inner skull surface. The noise covariance matrix was estimated from the prestimulus interval (−200 to 0 ms relative to stimulus onset). The same noise covariance matrix and inverse operator were used for both the attended and passive sessions.

Only responses to standard stimuli without gap were analyzed. For each epoch, source estimates were computed using the *apply_inverse_epochs* function from MNE-Python, enabling single-trial analysis in source space.

Further analysis focused on the superior temporal gyrus region (STG+) which comprised areas A1, 52, LBelt, PBelt, MBelt, RI, A4, TA2, PI, and PoI1 according to the HCPMMP1 atlas (Glasser, et al., 2016) (Figure 2). All these regions are structurally interconnected (Baker, et al., 2018; Rolls, Rauschecker, Deco, Huang, & Feng, 2023) and are involved in auditory information processing (Blenkmann, et al., 2019; Remedios, Logothetis, & Kayser, 2009; Tamura, Kuriki, & Nakano, 2015).

To define individual ROIs, the data were averaged across four stimulus types and two experimental conditions, and the orientation of source currents within the STG+ region was aligned using the ‘label_sign_flip’ function from MNE-Python. For each subject separately, we defined two ROIs in each hemisphere corresponding to (1) 30 vertices with maximal amplitude in 87–127 ms window (±20 ms around the N100m group maximum), and (2) 30 vertices with maximal amplitude in the later portion of the SW (300-600 ms), which did not overlap with transient components of the auditory response.

The rationale for the separate ‘N100m’ and ‘late SN’ ROIs was supported by statistical analysis. A paired Hotelling’s T² test revealed a significant difference between three-dimensional MNI-space-projected coordinates of the ‘N100m’ and ‘later SN’ maxima in the right hemisphere (F(3, 27) = 5.07, p = 0.006) and a trend for their difference in the left hemisphere (F(3, 27) = 2.65, p = 0.069) (Figure 2). N100m ROI was then used to analyse activity in 50-150 ms range, while the ‘later SN’ ROI – for the later time ranges.

To account for task-related factor (distance from the previous target, new target probability/expectancy), in the gap-detection task, epochs were grouped according to their position relative to the preceding target stimulus (see Results section for details). No such grouping was performed for the passive condition, where target stimuli were absent.

Next, time courses were extracted for each epoch within the ROIs using an orientation-constrained averaging procedure. Based on these time series, mean amplitudes were calculated within three time windows: 70–150 ms (in ‘N100m’ ROI) and 150–250 ms, 250–600 ms (in late SN ROI).

### 2.6 Statistical analysis

Statistical analyses were performed in R (Version 4.5.3).

To assess differences in omission errors across the four target carriers we used chi-square tests of independence. Following a significant overall effect, pairwise post-hoc chi-square tests compared each test condition (periodic non-vowels, non-periodic vowels, periodic vowels) to the control non-periodic non-vowel condition. Bonferroni correction with a corrected significance threshold of α = 0.0167 (0.05/3) was applied to account for multiple comparisons.

To investigate the relationship between omission errors and the order of the target trial relative to a previous target, we employed a generalized linear mixed-effects model (GLMM) with a binomial error distribution and logit link function using the glmer function from the lme4 package. The model included Order as a fixed effect and random intercepts for each subject to account for repeated measures within participants. The dependent variable was a binary indicator (1 = omission error, 0 = all other trial types). The statistical significance of fixed effects was assessed using Wald z-tests. Odds ratios (OR) were calculated by exponentiating the coefficient estimates.

To analyze reaction time (RT) we applied linear mixed-effects models (LMM) using the lme4 package. P-values were obtained using Satterthwaite’s approximation for degrees of freedom as implemented in the lmerTest package. The model included Order (trial order number following the previous target, treated as a continuous variable) and target Carrier (categorical variable with four levels corresponding to four stimulus types) as fixed effects. Random intercepts and random slopes for Order were included per subject to account for individual differences in baseline reaction times and order effects. The model was fitted using the bobyqa optimizer to ensure convergence.

All analyses of variance (ANOVAs) were conducted using the aov_ez function from afex package in R. To analyze effects of Trial Position (1^st^, 2^nd^, 3-4^th^, >4^th^) or Experiment (passive vs task) on the auditory MEG responses, three-way repeated-measures ANOVAs were conducted. The factors were Trial Position (or Experiment), Stimulus Type (4 levels), and Hemisphere (left vs right). The dependent variables were the differences in signal amplitude either between experimental conditions (task minus passive) or between stimuli types (each of test stimuli – periodic vowel, non-periodic vowel or periodic non-vowel minus control non-periodic non-vowel). These differences were averaged across Trial Positions. The mean number of averaged trials per subject and stimulus type was 22.2 for the 1^st^ after target Trial Position, 22.1 for the 2^nd^ group, 41.2 for 3-4^th^ group and 46.9 for the group with trial order numbers exceeding 4. Greenhouse-Geisser correction was applied to adjust for violations of sphericity assumption where appropriate. The corrected degrees of freedom are reported. Effect sizes are presented as generalized eta-squared (ges). Estimated marginal means were computed using the emmeans package in R. Pairwise comparisons were adjusted for multiple testing using the Tukey method (for comparisons among all levels of a factor) or Holm correction (for planned comparisons), as implemented in the emmeans function. All reported p-values are adjusted for multiple comparisons.

## 3 RESULTS

### 3.1 Behavioral results

Across subjects, the percentage of commission errors was negligibly low (<1% on average). The percentage of omission errors (i.e., no response or RT > 1s) is shown in figure 4A as a function of after-target trial number (left) and stimulus type (right).

**Figure 4.**
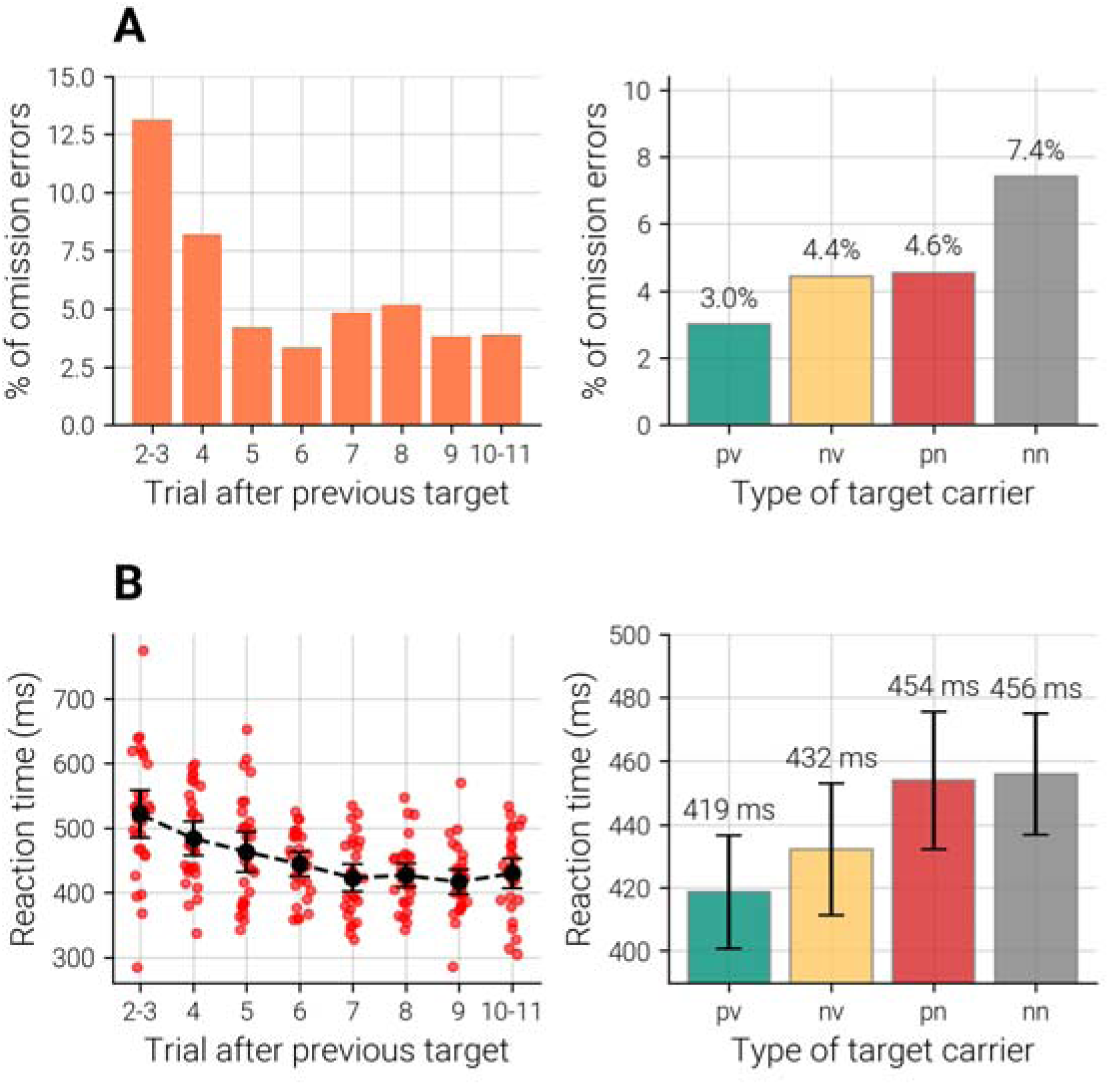
Behavioral results. (A) Percent of omission errors as a function of after-target trial number (left panel) and target carrier type (right panel). (B) Reaction time as a function of after-target trial (left panel) and target carrier type (right panel). Whiskers represent 95% confidence intervals. pv – periodic vowels, nv – non-periodic vowels, pn – periodic non-vowels, nn – non-periodic non-vowels (control stimulus).

A generalized linear mixed-effects model (binomial distribution, logit link; miss ∼ Order + (1 | subject) ) with Order as a categorical fixed effect (reference: order 2-3), revealed a significant main effect of trial Order on the likelihood of misses (b = −0.184, SE = 0.052, z = −3.53, p < .001). Miss rates were significantly lower than the reference value starting from order 5 (OR ≥ 0.22, p < 0.01 for all Orders ≥ 5). No significant difference was observed for order 4 (OR = 0.67, p = .330). Random intercept variance across subjects was 0.48 (SD = 0.70), indicating moderate individual differences in participants’ overall tendency to miss targets. These findings provide evidence that targets appearing closer to the previous target were more likely to result in omission compared to the later-appearing targets.

Chi-square tests of independence revealed a significant association between target carrier and miss rate (χ²(3) = 21.30, p < 0.001). The highest miss rate was observed for non-periodic non-vowel stimuli (∼7%), followed by non-periodic vowels (3.6%), periodic non-vowels (3.4%), and periodic vowels (2.2%). Post-hoc pairwise comparisons with Bonferroni correction (α = 0.0167) showed that all three experimental conditions had significantly lower miss rates than the non-periodic non-vowel baseline: nv (χ²(1) = 7.92, p_corrected = 0.015), pn (χ²(1) = 8.71, p_corrected = 0.009), and pv (χ²(1) = 16.31, p_corrected < 0.001).

RT decreased up to the seventh trial and then plateaued (Figure 4B). To test the statistical significance of this effect, we used linear mixed-effects model: RT ∼ Order + Carrier + (1 + Order | subject), where the Order corresponded to trials grouped according to their order number relative to the preceding target trial ( <=3, 4, 5, 6, >=7). Order was treated as a continuous variable, and target Carrier was a categorical variable with four levels. There was a significant negative linear effect of Order (F(1, 29.66) = 39.60, p < .001), with each consecutive Trial Position reducing RT by 21.75 ms per unit increase in trial Order (b = −21.75, t(29.66) = −6.29, p < .001). Carrier type also significantly affected RTs (F(3, 2523.83) = 13.72, p < .001), with responses being significantly faster when the gap occurred within periodic vowels (b = −35.26, t(2530.23) = −5.39, p < .001) or non-periodic vowels (b = −20.49, t(2519.64) = −3.22, p = .001) compared to the non-periodic non-vowel reference, while the periodic non-vowel stimuli did not differ significantly (b = −0.46, t(2523.89) = −0.07, p = .943). The full model explained 20% of the variance, with random effects accounting for 15%. A strong negative correlation (r = −0.93) was found between intercepts and slopes, suggesting that slower participants showed a greater decrease in reaction time with increasing Order.

To summarize, target expectancy increased as a function of the number of trials elapsed since the previous target and was accompanied by behavioral signatures of enhanced sustained attention. Specifically, omission errors decreased and reaction times shortened as target probability increased, consistent with progressively greater attentional engagement prior to target occurrence. Gap detection was most difficult when the target was embedded in the aperiodic non-vowel stimulus, as evidenced by the highest omission rate and the longest reaction times.

### 3.2 Effect of attention on the auditory cortical responses at different levels of target expectancy

Since both the probability of target occurrence and behavioral indices of attentiveness (error rate, RT) varied significantly as a function of the trial position following the target, we subdivided the trials into four groups to estimate the effect of attention:

1. *First trial after target*: Characterized by a prolonged interval following the previous (target) trial (see Figure 1) and a zero probability of target occurrence (89 trials per subject on average).
2. *Second trial after target*: Presented after a regular ISIs (500-800 ms), and associated with a vanishingly small probability of target occurrence (88.5 trials per subject on average).
3. *Trials 3-4*: Presented with regular ISIs (500-800 ms), associated with a low probability of target occurrence, and characterized by relatively longer RTs (164.7 trials per subject on average).
4. *Trials >4*: Presented with regular ISIs (500–800 ms), associated with a high likelihood of target occurrence, and characterized by relatively shorter RTs (187.5 trials per subject on average).

Figure 5 shows the difference in auditory-cortical evoked current time courses between the passive listening and gap-detection conditions. Relative to passive listening, directing attention to the auditory task enhanced an early negative response (∼50–150 ms) across all stimulus types and trial positions. This task-related enhancement is consistent with increased stimulus-driven (exogenous) attentional orienting to auditory events during active monitoring (Hillyard, Hink, Schwent, & Picton, 1973).

**Figure 5.**
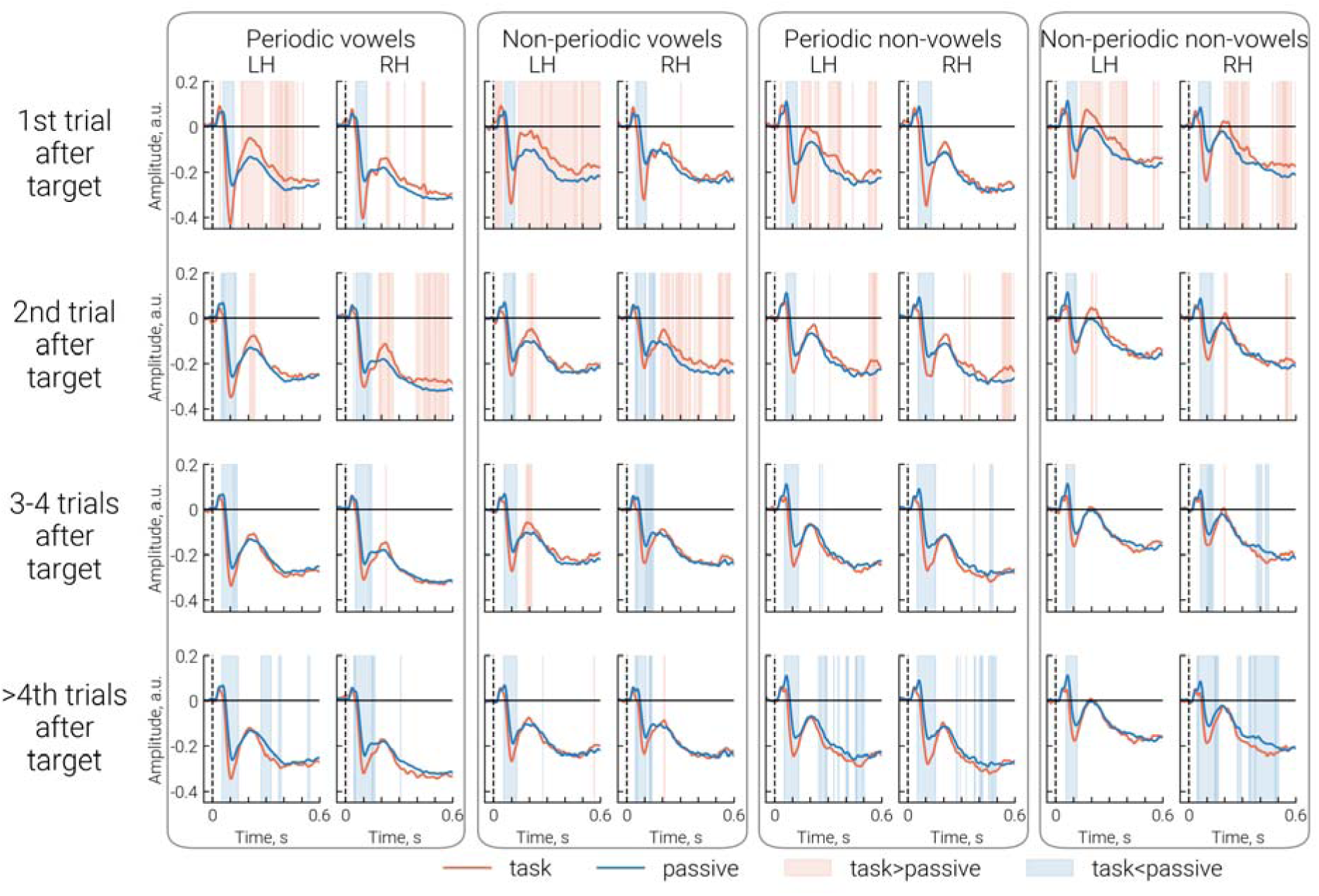
Grand-averaged neuronal current time courses during passive listening and the gap detection task. The four columns correspond to four stimuli types, and the four rows correspond to trial position relative to the preceding target event in the gap detection task (1st, 2nd, 3rd–4th, >4th). The *red line* shows the average time courses for non-target trials in the gap detection task. The *blue line* represents the time course averaged over all trials of each respective stimulus type in the passive listening condition and is replicated across all four rows within each column. LH and RH denote the left and right hemispheres, respectively. Blue and pink shaded areas indicate significant opposite-direction differences between the passive listening and gap detection conditions (p < 0.05, FDR-corrected).

To statistically assess how task-related effects depended on trial position following the target, stimulus type, and hemisphere, we performed repeated-measures ANOVAs. Within-subject factors included Trial Position (1st, 2nd, 3rd–4th, >4th trial relative to the preceding target event), Stimulus Type (four levels), and Hemisphere (left, right). The dependent variable was the difference in signal amplitude time courses between the task and passive listening conditions (Task minus Passive), averaged respectively for the 1st, 2nd, 3rd–4th, and >4th post-target trials.

The analysis was conducted separately for three time intervals assumed to reflect partly distinct neural processes. The first interval (70–150 ms; ‘early SN interval’) overlaps with processing negativity (PN) and the auditory N100m response and is associated with early sensory encoding and automatic attention orienting (Näätänen, Kujala, & Winkler, 2011; Rosburg, Boutros, & Ford, 2008). The second interval (150–250 ms; ‘middle SN interval’) includes the P200m component (Crowley & Colrain, 2004) and encompasses later-stage sensory encoding and stimulus classification processes that are sensitive to perceptual learning, training, and familiarity (Bidelman, Moreno, & Alain, 2013; Crowley & Colrain, 2004; Ross & Tremblay, 2009; Shahin, Roberts, Pantev, Trainor, & Ross, 2005; Steinmetzger & Rupp, 2024; Tremblay, Ross, Inoue, McClannahan, & Collet, 2014). The third interval (250–600 ms; ‘late SN’ interval) corresponds to the later portion of the sustained negativity that does not overlap with transient auditory responses and reflects sustained representation of the auditory object (Gutschalk & Uppenkamp, 2011). Table 1 shows the ANOVA results for these three intervals.

**Table 1.** Repeated-measures ANOVA results for the task-related modulation of auditory cortex responses expressed as the difference between the gap-detection and passive listening conditions within the early, middle, and late response intervals.

| Effect | Df | F | ges | p-value |
| --- | --- | --- | --- | --- |
| **Early interval: 70-150 ms** |  |  |  |  |
| **Trial Position<sup>#</sup>** | **1.99, 57.83** | **11.47 \*\*\*** | **.043** | **<.001** |
| **Stimulus Type** | **2.76, 80.17** | **9.16 \*\*\*** | **.035** | **<.001** |
| Hemisphere | 1, 29 | 0.11 | <.001 | .746 |
| Trial Position x Stimulus Type | 5.96, 172.79 | 0.61 | .003 | .722 |
| Trial Position x Hemi | 1.96, 56.76 | 1.44 | .003 | .245 |
| Stimulus Type x Hemi | 2.78, 80.76 | 1.08 | .003 | .359 |
| Trial Position x Stimulus Type x Hemi | 6.64, 192.44 | 0.92 | .003 | .487 |
| **Middle interval: 150-250 ms** |  |  |  |  |
| **Trial Position** | **1.66, 48.22** | **12.22 \*\*\*** | **.057** | **<.001** |
| Stimulus Type | 2.73, 79.25 | 1.45 | .005 | .236 |
| Hemisphere | 1, 29 | 1.75 | .010 | .196 |
| Trial Position x Stimulus Type | 5.63, 163.26 | 0.54 | .002 | .771 |
| **Trial Position x Hemi** | **1.89, 54.75** | **5.53 \*\*** | **.009** | **.007** |
| Stimulus Type x Hemi | 2.84, 82.30 | 0.29 | <.001 | .820 |
| Trial Group x Stimulus Type x Hemi | 6.68, 193.78 | 0.70 | .002 | .663 |
| **Late interval: 250-600 ms** |  |  |  |  |
| **Trial Position** | **2.24, 64.82** | **22.57 \*\*\*** | **.070** | **<.001** |
| Stimulus Type | 2.66, 77.23 | 1.50 | .008 | .225 |
| Hemisphere | 1, 29 | 0.07 | <.001 | .788 |
| Trial Position x Stimulus Type | 6.65, 192.95 | 1.25 | .006 | .280 |
| Trial Position x Hemi | 2.39, 69.41 | 2.32 | .006 | .096 |
| Stimulus Type x Hemi | 2.35, 68.07 | 0.24 | <.001 | .817 |
| Trial Position x Stimulus Type x Hemi | 6.32, 183.26 | 1.63 | .006 | .136 |

Note: dependent variable was the difference in auditory response time courses between the gap detection task and the passive condition (Task minus Passive).
Here and hereafter Greenhouse-Geisser corrected degrees of freedom are reported when applicable; ges – generalized eta-squared. \*\* $p < 0.01$ , \*\*\* $p < 0.001$
<sup>#</sup> Trial Position after target (1<sup>st</sup>, 2<sup>nd</sup>, 3-4<sup>th</sup>, >4<sup>th</sup>).

The factor of Trial Position was significant for all time intervals (Figure 6).

**Figure 6.**
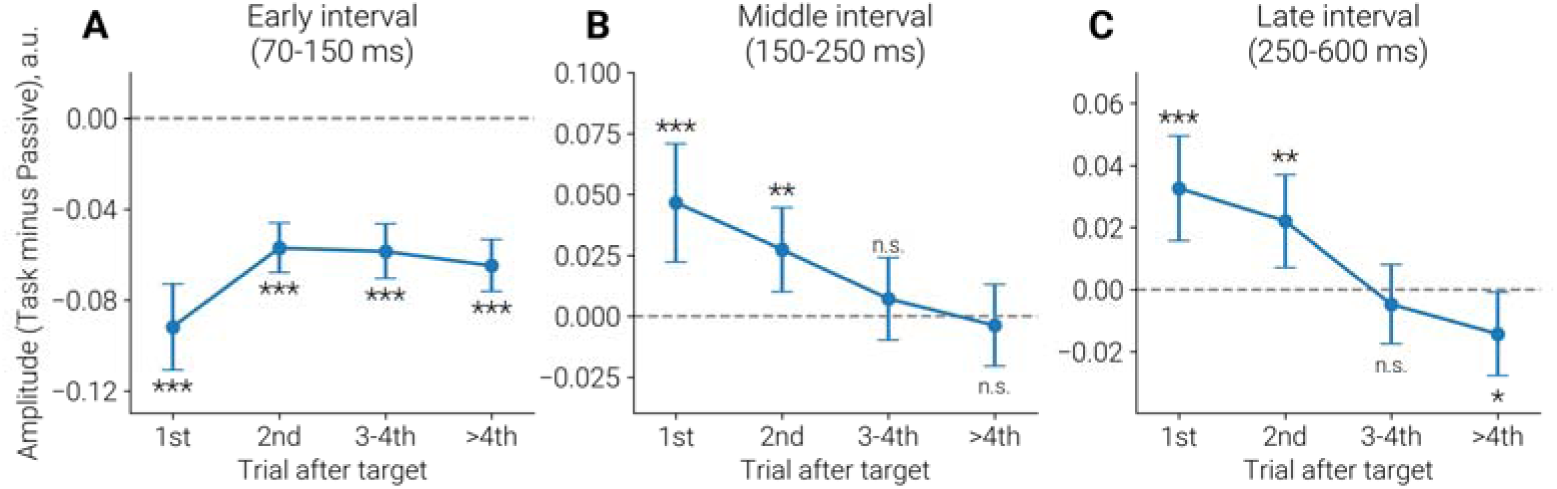
Effects of trial position relative to the preceding target event on task-related modulation of auditory cortical evoked responses within the early (A), middle (B), and late (C) response intervals. Task-related modulation was quantified as the difference in mean response amplitude between the task and passive listening conditions (Task minus Passive) and is plotted as a function of post-target trial position. Dashed lines indicate the absence of a difference between conditions (i.e., Task – Passive = 0). Negative values correspond to greater negativity during the Task than during Passive listening. *P*-values indicate the probability of a non-zero Task – Passive amplitude difference. * p < 0.05, ** p < 0.01, *** p < 0.001

During the early SN interval, negativity increased in the auditory task relative to passive listening across all Trial Positions (Figure 6A). Pairwise comparisons of estimated marginal means showed that task-related negativity in the first trial following the target was significantly greater than in all subsequent Trial Positions (2nd, 3rd–4th, >4th; all Tukey-adjusted p ≤ .019), whereas no significant differences were observed among the latter three groups (all p ≥ .46). A strong main effect of Stimulus Type was also present. Tukey-adjusted pairwise comparisons revealed that, during this early time interval, the task-related increase in negativity for periodic non-vowels was greater than for all other stimuli types (non-periodic non-vowels: p = .003, non-periodic vowels: p < .001, periodic vowels: p = .07) (*Supplementary figure S1*).

The middle SN interval was characterized by a task-related reduction in negativity (i.e., increased positivity) during the first two post-target trials, with little or no task-related modulation in subsequent trials (Figure 6B). A significant Trial Position × Hemisphere interaction reflected a greater decrease in task-related negativity in the left hemisphere during the first post-target trial (*Supplementary figure S2*).

In the late SN interval, negativity was likewise reduced during the task relative to passive listening in the first two post-target trials, but increased in later trials (>4th), exceeding the level observed in the passive condition (Figure 6C).

To summarize, the auditory task modulated auditory-cortex responses differently across the early, middle, and late SN intervals. In the early interval, task-related negativity was enhanced across all post-target trial positions, with the largest effect observed immediately after a target. In contrast, the middle and late intervals showed reduced negativity during the first post-target trials. This reduction gradually diminished as the number of trials since the preceding target increased, eventually giving way to enhanced negativity relative to passive listening in later trials, consistent with increasing target expectancy.

### 3.3 Effect of attention on sustained processing negativity (SPN)

The effect of sustained attention on SPN—a differential response reflecting SN enhancement associated with the processing of acoustic regularities characteristic of vowels—is of particular interest in the present study. Reduced attentional engagement with speech sounds may contribute to the atypical neural responsiveness to speech observed in clinical populations with neurodevelopmental disorders. Therefore, we next assessed the effect of task on the SPN, defined as the differential auditory-cortex response to stimuli containing temporal and/or spectral regularity (periodic vowels, periodic non-vowels, and non-periodic vowels) relative to aperiodic noise. Separate ANOVAs were conducted for the early, middle, and late response intervals, with SPN amplitude as the dependent variable and Experiment (passive listening vs. gap-detection task), Stimulus Type, and Hemisphere as within-subject factors.

#### 3.3.1 SPN in the early interval (70-150 ms)

Figure 7 illustrates the SPN in the auditory cortical region with maximal N100m negativity, focusing on the early interval. Because the previous analysis revealed pronounced task-related differences in early-interval responses between the first post-target trial and subsequent post-target trials (Figure 6A), we conducted separate repeated-measures ANOVAs for the first versus later post-target trials (Table 2).

**Figure 7.**
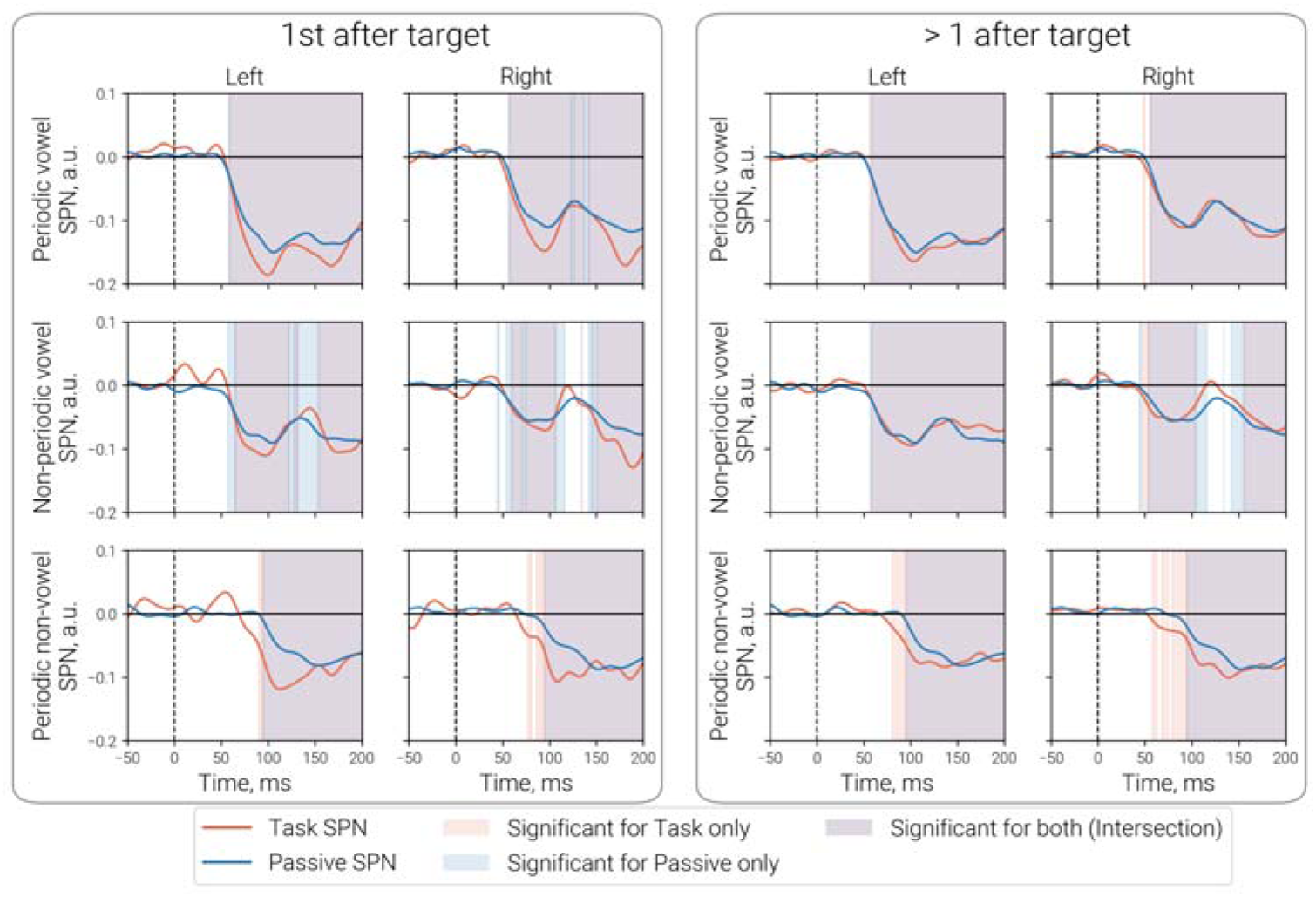
Time course of sustained processing negativity (SPN) in the N100m cortical ROI. SPN was computed as the difference between responses to test stimuli containing acoustic regularity and control stimuli containing acoustic noise (Test minus Control). The timescale is limited to 200 ms to highlight the early (N100m) response interval. Blue and red lines denote passive and task conditions, respectively. Pink shading indicates significant negative SPN during task performance; blue shading indicates significantly negative SPN in the passive condition; purple shading indicates overlap between the two conditions. Note the earlier onset of SPN for periodic non-vowels during attention to the auditory stream relative to passive listening. Probability values are FDR-corrected (*p* < 0.05).

**Table 2.** Repeated-measures ANOVA results for the SPN (difference in responses between Test and Control stimuli) in the early response interval (70-150 ms).

| Effect | df | F | ges | p-value |
| --- | --- | --- | --- | --- |
| **1<sup>st</sup> post-target trial** |  |  |  |  |
| Experiment <sup>#</sup> | 1, 29 | 5.21* | .023 | .03 |
| Stimulus Type | 1.43, 41.39 | 36.32 *** | .145 | <.001 |
| Hemisphere | 1, 29 | 4.88* | .031 | .035 |
| Experiment x Stimulus Type | 1.92, 55.82 | 4.94* | .011 | .011 |
| Experiment x Hemi | 1, 29 | 0.28 | <.001 | .601 |
| Stimulus Type x Hemi | 1.83, 53.15 | 4.49* | .013 | .018 |
| Experiment x Stimulus Type x Hemi | 1.96, 56.96 | 0.03 | <.001 | .967 |
| **>1<sup>st</sup> post-target trials** |  |  |  |  |
| Experiment <sup>#</sup> | 1, 29 | 1.41 | .005 | .245 |
| Stimulus Type | 1.49, 43.29 | 62.75 *** | .208 | <.001 |
| Hemisphere | 1, 29 | 7.92** | .051 | .009 |
| Experiment x Stimulus Type | 1.96, 56.85 | 12.67 *** | .012 | <.001 |
| Experiment x Hemi | 1, 29 | 0.88 | .002 | .354 |
| Stimulus Type x Hemi | 1.55, 45.80 | 9.10 ** | .035 | .001 |
| Experiment x Stimulus Type x Hemi | 1.90, 55.03 | 1.85 | .002 | .168 |

\* $p < 0.05$ , \*\* $p < 0.01$ , \*\*\* $p < 0.001$

A significant main effect of Stimulus Type was observed for both the first after-target trial and all consecutive trials, indicating that early SPN was largest for the “most regular” periodic vowel stimuli compared with the other test stimuli (periodic non-vowels and non-periodic vowels), replicating the previous findings obtained in independent samples (Fadeev, et al., 2024; Gutschalk & Uppenkamp, 2011; Orekhova, et al., 2024). The significant Stimulus Type × Hemisphere interactions indicated a left-hemispheric predominance for vowel stimuli, both periodic and non-periodic (Holm-adjusted ps ≤ .027), whereas no such lateralization was observed for periodic non-vowels (p ≥.51). This left-hemispheric predominance of the SPN for vowels also replicates previous findings and is discussed in details in (Orekhova, et al., 2024).

A significant main effect of Experiment was also observed for both the first and subsequent post-target trials. However, this main effect was accounted for by the Experiment × Stimulus Type interaction (Figure 8). Holm-adjusted pairwise comparisons of estimated marginal means showed that the task-versus-passive contrast was significant only for periodic non-vowels (ps < .005 in both trial positions), but not for periodic or non-periodic vowels (all ps > .1). As illustrated in Figure 5, sustained attention during the gap-detection task led to an earlier onset and greater amplitude of the SPN for periodic non-vowel stimuli during the early interval.

**Figure 8.**
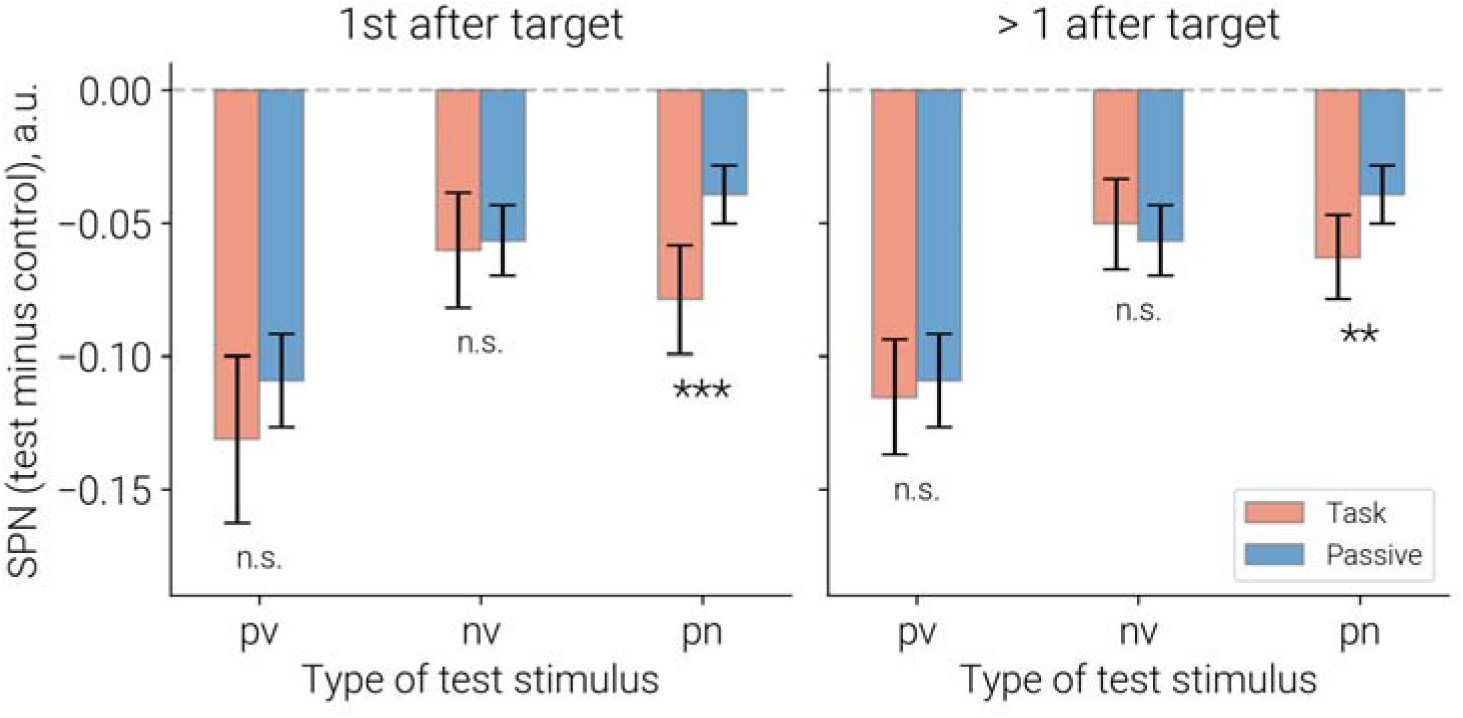
Experiment × Stimulus Type interaction for the sustained processing negativity (SPN: Test minus Control) in the early (70-150 ms) interval. (**A)** First trial following the target; (**B**) trials >1 following the target. pv – periodic vowels, nv – non-periodic vowels, pn – periodic non-vowels. ** p < 0.01, *** p < 0.001

#### 3.3.2 SPN in the middle (150-250 ms) and late (250-600 ms) intervals

Figure 9 illustrates SPN in the left and right cortical ROIs with maximal SN amplitude in the late interval (‘late SN’ ROIs). The previous analysis has shown that in these intervals the SN was strongly modulated by proximity to the preceding target. During the first two post-target trials, SN was reduced during the auditory task relative to passive listening. In subsequent trials, task-related differences were either absent or reversed, with SN becoming enhanced during the auditory task (Figure 6B,C). Accordingly, we conducted separate ANOVAs for two groups of post-target trials: trials 1–2 and trials >2 following the target (Tables 3 and 4).

**Figure 9.**
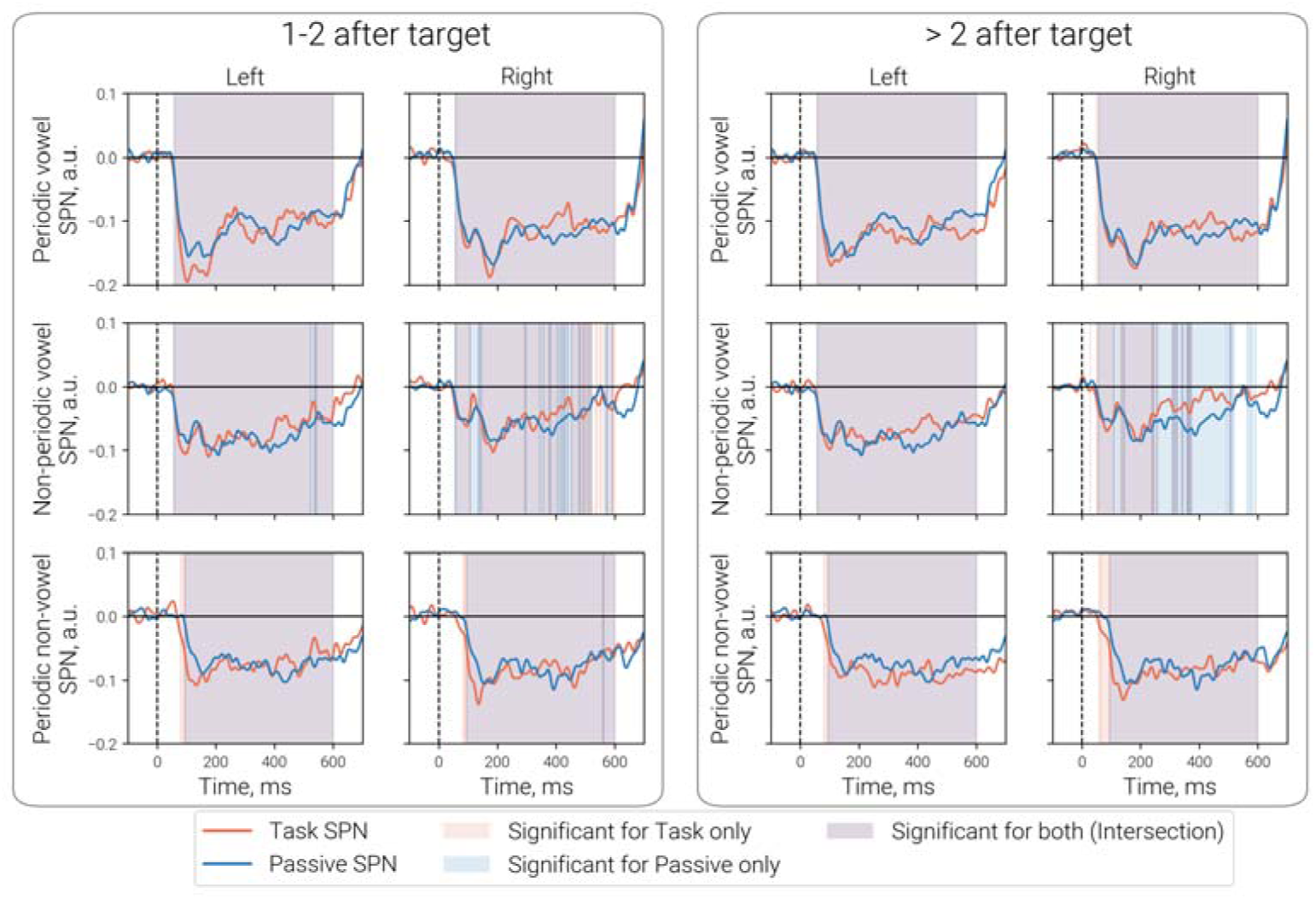
Time course of sustained processing negativity (SPN: Test minus Control) in the ‘late SN’ ROIs. (**A**) First two trials following the target. (**B**) All trials excluding first and second post-target trials. Pink shading indicates significant negative SPN during task performance; blue shading indicates significantly negative SPN in the passive condition, and purple shading indicates overlap between these conditions. Probability values are FDR-corrected (p < 0.05).

**Table 3.** Repeated-measures ANOVA results for the SPN (test conditions – control condition) in the middle interval (150-250 ms).

| Effect | df | F | ges | p-value |
| --- | --- | --- | --- | --- |
| **1-2<sup>nd</sup> post-target trial** |  |  |  |  |
| Experiment<sup>#</sup> | 1, 29 | 0.16 | <.001 | .688 |
| **Stimulus Type** | **1.97, 57.20** | **30.10 \*\*\*** | **.141** | **<.001** |
| Hemisphere | 1, 29 | 0.14 | <.001 | .709 |
| Experiment x Stimulus Type | 1.79, 51.78 | 0.45 | <.001 | .621 |
| Experiment x Hemi | 1, 29 | 0.07 | <.001 | .797 |
| Stimulus Type x Hemi | 1.67, 48.36 | 3.36 | .009 | .051 |
| Experiment x Stimulus Type x Hemi | 1.99, 57.67 | 0.45 | <.001 | .640 |
| **> 2<sup>nd</sup> post-target trials** |  |  |  |  |
| Experiment <sup>#</sup> | 1, 29 | 0.02 | <.001 | .882 |
| **Stimulus Type** | **1.63, 47.19** | **31.80 \*\*\*** | **.159** | **<.001** |
| Hemisphere | 1, 29 | 0.06 | <.001 | .807 |
| **Experiment x Stimulus Type** | **1.92, 55.61** | **4.53 \*** | **.007** | **.016** |
| Experiment x Hemi | 1, 29 | 0.01 | <.001 | .944 |
| **Stimulus Type x Hemi** | **1.73, 50.27** | **4.40 \*** | **.014** | **.022** |
| Experiment x Stimulus Type x Hemi | 1.96, 56.86 | 0.18 | <.001 | .831 |

**Table 4.** Repeated-measures ANOVA results for the SPN (test conditions minus control condition) in the late interval (250-600 ms).

| Effect | df | F | ges | p-value |
| --- | --- | --- | --- | --- |
| **1-2<sup>nd</sup> post-target trial** |  |  |  |  |
| Experiment <sup>#</sup> | 1, 29 | 0.31 | .001 | .580 |
| **Stimulus Type** | **1.92, 55.74** | **18.32 \*\*\*** | **.100** | **<.001** |
| Hemisphere | 1, 29 | 0.33 | .002 | .572 |
| Experiment x Stimulus Type | 1.62, 47.01 | 0.11 | <.001 | .854 |
| Experiment x Hemi | 1, 29 | 0.02 | <.001 | .890 |
| **Stimulus Type x Hemi** | **1.92, 55.72** | **6.17 \*\*** | **.015** | **.004** |
| Experiment x Stimulus Type x Hemi | 1.86, 54.06 | 0.34 | <.001 | .698 |
| **> 2<sup>nd</sup> post-target trials** |  |  |  |  |
| Experiment <sup>#</sup> | 1, 29 | 0.06 | <.001 | .813 |
| **Stimulus Type** | **1.97, 57.02** | **31.81 \*\*\*** | **.152** | **<.001** |
| Hemisphere | 1, 29 | 1.98 | .012 | .170 |
| **Experiment x Stimulus Type** | **1.96, 56.79** | **5.93 \*** | **.010** | **.005** |
| Experiment x Hemi | 1, 29 | 1.62 | .003 | .213 |
| **Stimulus Type x Hemi** | **1.94, 56.29** | **8.46 \*\*\*** | **.020** | **<.001** |
| Experiment x Stimulus Type x Hemi | 1.97, 57.27 | 0.05 | <.001 | .949 |

\* p < 0.05, \*\* p < 0.01, \*\*\* p < 0.001

Similar to the early interval, significant main effects of Stimulus Type were observed in both the middle and late intervals, irrespective of trial position and experiment (passive listening vs. auditory task). These effects reflected larger SPN amplitudes for periodic vowels than for stimuli lacking either temporal periodicity or formant structure (i.e., periodic non-vowels and non-periodic vowels).

A significant Stimulus Type × Hemisphere interaction was found for later post-target trials (>2 after the target) in the middle interval and for both trial positions in the late interval. However, Holm-corrected pairwise comparisons of the signal amplitude in the left and right hemispheres revealed significant differences only for the late SPN: non-periodic vowels elicited greater negativity in the left hemisphere (ps ≤ .002), whereas periodic stimuli showed no significant lateralization (all ps ≥ .49). These results are generally in agreement with previous findings of greater left-hemispheric lateralization for processing of formant structure of vowels (Orekhova, et al., 2024).

A significant Experiment × Stimulus Type interaction was observed in both the middle and late intervals, but only for trials occurring more than two trials after the preceding target (Figure 8). This interaction was primarily driven by a task-related reduction in SPN elicited by the non-periodic vowels. Holm-corrected pairwise comparisons confirmed that this reduction reached significance in the late SPN interval (p = .047).

To summarize, task-related attentional engagement exerted stimulus-specific effects on SPN that differed markedly across temporal intervals. In the early interval (70–150 ms), the auditory task selectively enhanced SPN for periodic non-speech stimuli, leading to an earlier onset and higher amplitude of the response relative to passive listening. In contrast, the late SPN interval (250-600 ms) showed selective sensitivity to non-periodic vowels: the task was associated with a reduction of SPN for these stimuli, whereas no comparable modulation was observed either for the more natural periodic vowels or for non-speech periodic sounds.

## 4 DISCUSSION

In this study, we examined whether fluctuations in sustained attention and target expectancy during a gap-detection task modulate the processing advantage of acoustically regular sounds—vowels and non-vocal temporally regular sounds—relative to spectrally matched noise. Passive listening during salient video viewing served as a control condition.

Processing of prolonged spectrally complex sounds is associated with an early-onset, long-lasting negativity – SN – that overlaps with transient auditory responses (Fadeev, et al., 2026; Gutschalk, Patterson, Scherg, Uppenkamp, & Rupp, 2004; Orekhova, et al., 2024). The SN is enhanced for acoustically regular sounds relative to spectrally matched noise (Barascud, Pearce, Griffiths, Friston, & Chait, 2016; Sohoglu & Chait, 2016; Southwell, et al., 2017). Here, we quantified this differential response—termed sustained processing negativity (SPN)—as a neural index of enhanced auditory processing. Importantly, in the gap-detection task, all sounds could potentially contain the target stimulus (a temporal gap), requiring sustained attention until stimulus offset. Consequently, all non-target trials remained behaviourally relevant throughout the experiment.

We found that both decreases and increases in sustained attention—induced, respectively, by attention diversion following rare targets and by enhanced engagement associated with rising target expectancy—significantly modulated auditory responses across all stimulus types. However, these attentional fluctuations did not alter the SPN associated with periodic vowels. In contrast, for non-vocal periodic acoustic patterns, task-related orienting to stimulus onset led to an earlier emergence of the SPN, suggesting facilitated discrimination of temporal regularity.

Given the limited evidence directly comparing the effects of task-driven orienting and non-selective attention on auditory sustained responses, we consider these mechanisms jointly in relation to our central finding: namely, the relative stability of the vowel processing advantage.

### 4.1 Sensory orienting to stimulus onset enhances the early sustained negativity

The neural response observed approximately 70–150 ms after stimulus onset (‘N100m interval’) has been linked to sensory orienting to auditory events (Hillyard, Hink, Schwent, & Picton, 1973; Näätänen, Kujala, & Winkler, 2011). Consistent with previous literature (Alho, Sams, Paavilainen, Reinikainen, & Näätänen, 1989; Hansen & Hillyard, 1980; Raudonat, Doroszewski, & Gutschalk, 2025), we observed increased negativity in N100m interval during task performance relative to passive listening (Figure 6A). This increase was greatest for the first stimulus following a target and subsequently stabilized across later trials.

The pronounced enhancement of early negativity in the first post-target trial may reflect a transient disengagement from the ongoing task following target detection, rendering the subsequent stimulus relatively unexpected and increasing the salience of its onset (Lavro, Ben-Shachar, Saville, Klein, & Berger, 2018). In addition to attentional reorienting, the prolonged interval preceding the first post-target stimulus may itself contribute to this effect, as N100/N100m amplitude is known to increase with longer interstimulus intervals(Campbell & Neuvonen, 2007; Rosburg & Mager, 2021).

In contrast to this transient post-target effect, the relatively uniform increase in early negativity across subsequent task trials (2nd, 3rd–4th, and >4th post-target trials) relative to passive listening may reflect a relatively sustained top-down attentional gating mechanism that enhances sensory processing of attended stimuli (Hillyard, Hink, Schwent, & Picton, 1973; Näätänen, Kujala, & Winkler, 2011).

### 4.2 Proximity to the previous target and target expectancy modulate later sustained negativity

The direction of changes in later SN intervals (150–250 ms and 250–600 ms) depended critically on two factors: proximity to the preceding target and expectancy of an upcoming target.

#### 4.2.1 Proximity to a previous target

In the later SN intervals, the effect of proximity to the preceding target was opposite to that observed during the early SN interval. Specifically, the amplitude of the later SN evoked by the first post-target stimulus was reduced relative to passive listening across all stimulus types (Figures 5, 6B,C). In the 150–250 ms interval, this reduction tended to be greater in the left hemisphere (*Supplementary Figure S2*). The reduction remained evident during the second post-target trial and vanished rapidly thereafter.

This paradoxical reduction of the later SN under attention, relative to passive listening, is most parsimoniously explained by a central-processing “bottleneck” effect induced by rare salient events (Lavro, Ben-Shachar, Saville, Klein, & Berger, 2018). According to this account, processing of salient and infrequent targets temporarily consumes limited cognitive resources, thereby constraining processing of subsequent stimuli. Such resource reallocation primarily affects later, higher-order stages of processing (Houtman & Notebaert, 2013; Jentzsch & Dudschig, 2009; Lavro, Ben-Shachar, Saville, Klein, & Berger, 2018; Notebaert, et al., 2009), which may explain the absence of this effect during the earlier SN interval. From this perspective, the opposite effects of a rare target on early and middle-late SN are consistent with a combination of enhanced sensory orienting at sound onset and reduced availability of central processing resources during later stages of auditory analysis.

#### 4.2.2 Target expectancy

In the gap-detection task, targets never occurred immediately after a preceding target and were extremely rare in the second post-target trial. Consequently, the probability of target occurrence increased progressively with the number of trials since the previous target, reaching its maximum by the sixth post-target trial (Figure 2A,B). This increase in target probability was accompanied by increasing target expectancy and may explain the corresponding improvement in behavioral performance, reflected in reduced omission errors after the third post-target trial and shorter reaction times to targets (Figure 4).

Accordingly, the enhancement of the late SN beyond the level observed during passive listening (Figure 6C) may reflect increasing expectancy of the target, which in turn enhances non-selective attention to the auditory stream independently of specific stimulus features. The absence of a comparable expectancy effect during the middle SN interval is less readily explained. One possibility is that this interval overlaps with the transient P200m component, whose amplitude is also sensitive to target probability (Fishman, Lee, & Sussman, 2021).

### 4.3 The processing advantage of periodic vowels over noise remains stable despite task-related modulation of sustained negativity

Consistent with previous literature (Barascud, Pearce, Griffiths, Friston, & Chait, 2016; Gutschalk, Patterson, Scherg, Uppenkamp, & Rupp, 2007; Hewson-Stoate, Schönwiesner, & Krumbholz, 2006; Southwell, et al., 2017; Southwell & Chait, 2018), we observed enhanced negativity in response to auditory patterns—periodic and aperiodic vowels, as well as periodic non-vocal sounds—relative to spectrally matched noise. This SPN emerged rapidly (50–70 ms after stimulus onset) and persisted throughout stimulus presentation during both passive listening and the gap-detection task (Figure 7).

During the gap detection task, attention-related processes—including early orienting, post-target attention diversion, and target expectancy—strongly modulated sustained auditory responses, often in opposite directions across different time intervals. These findings indicate considerable flexibility of the SN under top-down control and raise the question of whether such modulations also affect the processing advantage of structured auditory patterns over noise, as indexed by the SPN.

To address this question, we compared SPN across experimental conditions, analyzing the early and late response intervals separately. We further examined whether task effects varied as a function of proximity to the preceding target (first post-target trial vs. subsequent trials) and target expectancy (the first two post-target trials, associated with minimal target probability, versus the third and later post-target trials).

#### 4.3.1 Early SPN: orienting facilitates early processing of non-vocal periodic patterns but not vowels

Stimulus-driven orienting enhanced the early phase of SPN (70–150 ms) primarily for non-vocal periodic patterns, while leaving it largely unchanged for both vowel types—periodic and aperiodic (Figures 7 and 8).

Relative to vowel stimuli, the onset of the SPN for periodic non-vocal sounds was delayed by approximately 30 ms under passive listening conditions (≈90 ms for periodic non-vocal sounds vs. ≈60 ms for periodic and non-periodic vowels). In addition to increasing the magnitude of the SPN, task-related orienting also accelerated its onset for non-vocal periodic sounds. As a result, under task conditions, this delay was largely eliminated in the right hemisphere (SPN onset at 50–60 ms for all stimuli) and decreased by half in the left hemisphere (Figure 7).

Overall, these findings are consistent with previous evidence demonstrating that the auditory system can extract cues to periodicity from very brief sound segments (approximately 3–5 glottal pulses; (Yrttiaho, Tiitinen, Alku, Miettinen, & May, 2010)), corresponding to approximately 36–60 ms in the present study. Temporal regularities in continuous acoustic sequences can be detected even in the absence of directed attention, as reflected in enhanced sustained responses (Barascud, Pearce, Griffiths, Friston, & Chait, 2016; Southwell, et al., 2017; Southwell & Chait, 2018) and in the pitch/periodicity onset response (POR), a cortical response to the emergence of periodicity observed during passive listening (Kim, et al., 2022; Krumbholz, Patterson, Seither-Preisler, Lammertmann, & Lütkenhöner, 2003).

Within this framework, the early component of the SPN evoked by periodic non-vocal sounds observed in the present study may partly reflect POR-like mechanisms involved in the rapid extraction of temporal periodicity in the auditory cortex. Importantly, we extend previous findings by showing that task-related orienting to stimulus onset—despite the absence of any explicit requirement to detect temporal regularity—selectively facilitates the early emergence of SPN for non-vocal periodic sounds (Figure 5). This finding suggests that the neural detection of temporal regularity in non-vocal sounds benefits from transient attention orienting to auditory events. More broadly, these results inform ongoing debates regarding whether the bottom-up saliency of non-vocal temporal regularities is sufficient to capture attention (Southwell, et al., 2017). Our findings suggest that such regularities may enhance early stimulus-driven orienting at sound onset but do not appear to elicit enhanced processing at the later stages (>150 ms), as discussed below.

In contrast, neural discrimination of vowel spectral structure emerged substantially earlier than discrimination of periodicity in non-vocal sounds and showed little additional facilitation under the task compared with passive listening (Figures 7, 8). Thus, unlike non-vocal periodicity, whose processing was accelerated by task-related orienting, vowel-related processing appeared to be largely optimized already during passive listening. Vowels constitute highly familiar and biologically relevant acoustic patterns that convey important conspecific information, and their processing may therefore rely predominantly on intrinsic saliency. This intrinsic saliency, or “vowelness,” may also contribute to the more rapid detection of interruptions (gaps) within vowel sounds relative to non-vocal sounds, as observed in the present study (Figure 4B).

### 4.4 Late SPN: vowel advantage remains stable despite task-related fluctuations in sustained attention

We observed no effect of task versus passive listening on the SPN throughout the 150-250 and 250-600 ms intervals during the first two post-target trials (Tables 3 and 4). This suggests that target-induced attentional diversion, although reducing overall sustained negativity, does not alter the higher-order processing advantage of auditory patterns, which appears relatively insensitive to transient shifts of attention away from the ongoing perceptual task.

Target expectancy, gradually emerging from the third post-target trial onward, exerted more stimulus-specific effects on late SPN (Tables 3 and 4). It reduced SPN for non-periodic vowels while leaving SPN for periodic vowels and periodic non-vocal sounds largely unchanged (Figure 10).

**Figure 10.**
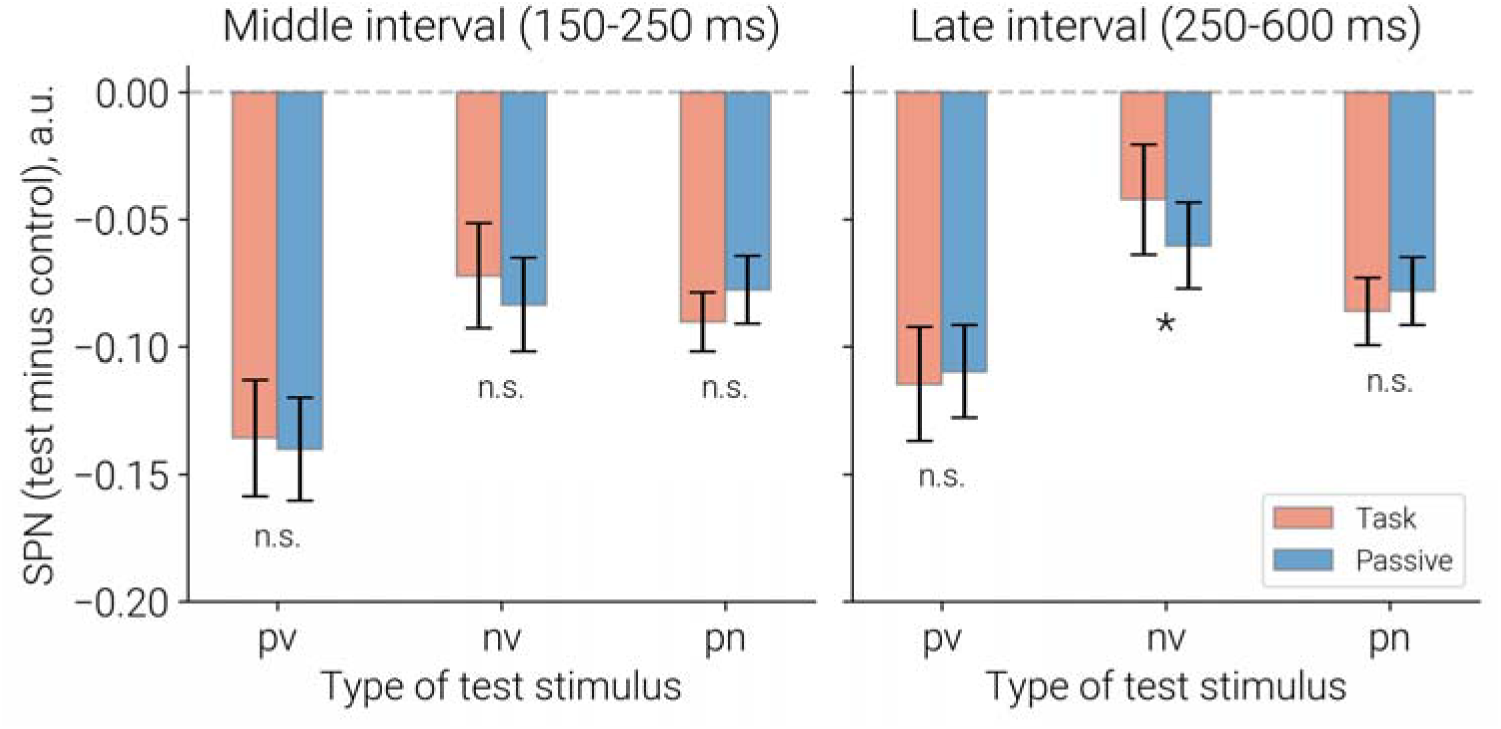
Experiment × Stimulus Type interactions in the middle (150-250 ms) (A) and late (250-600 ms) (B) SPN intervals. The plots show the effect of experimental condition on SPN amplitude for post-target trials occurring more than two trials after the preceding target (i.e., trials > 2). pv – periodic vowels, nv – non-periodic vowels, pn – periodic non-vowels. * p < 0.05

For non-periodic vowels, the task-related reduction in their advantage over noise may reflect the greater perceptual difficulty of detecting gaps within the control stimulus (aperiodic non-vocal noise). Consistent with this possibility, reaction times were longer for targets embedded in noise than in non-periodic vowels (456 vs. 432 ms). The increased difficulty associated with the noise carrier may have enhanced anticipatory monitoring of these stimuli, thereby contributing to the proportionally greater increase in late SN during later post-target trials (Figure 5) and, consequently, to the reduction of the SPN.

A similar account may explain the absence of task-related modulation of the SPN elicited by periodic non-vocal sounds. Gap detection in these stimuli was behaviorally as difficult as in aperiodic noise (mean RTs: 454 and 456 ms, respectively), suggesting a comparable degree of task-related monitoring. Consistent with this interpretation, periodic non-vocal sounds also exhibited expectancy-related increases in SN (Figure 5). As a result, both the periodic non-vocal sounds and the noise control showed similar task-related enhancements, leaving the differential SPN response largely unchanged.

In contrast, periodic vowels—arguably the stimuli most closely resembling natural speech sounds—elicited the largest SPN under both passive listening and task conditions (Figures 9 and 10). This advantage remained remarkably stable despite the modest task-related changes in SN observed for the control noise stimulus. Thus, the most parsimonious interpretation of the present findings is that target expectancy, and the associated increase in sustained attention to the auditory stream, has little influence on the intrinsic neural processing advantage conferred by the spectrotemporal regularities of periodic vowel sounds.

## 5 Conclusions

Our findings reveal dissociation between the effects of task-related orienting, post-target attentional diversion, and expectancy on neural processing of sounds characterized by periodicity or vowel formant structure.

Task-induced orienting to the auditory stream enhanced early neural negativity (70–150 ms) across all stimulus types and facilitated discrimination of periodicity in non-vocal sounds independently of target expectancy. In contrast, during late stage of processing (250–600 ms), sustained negativity tracked target expectancy, reflecting anticipatory monitoring of the auditory stream.

Importantly, these top-down modulations did not alter the intrinsic processing advantage of regular vowel sounds, which remained stable across task and passive listening conditions. Thus, although non-selective attention reshapes the temporal dynamics of sustained auditory responses, the neural prioritization of vowel spectrotemporal structure operates largely independently of such attentional fluctuations.

These findings contribute to ongoing debates concerning the extent to which intrinsically salient sound patterns capture attention (Southwell, et al., 2017), suggesting that vowel processing is supported by mechanisms that are relatively independent of attentional state. More broadly, the present results provide a framework for interpreting potential alterations in vowel processing in clinical populations characterized by deficits in sustained attention.

## Author Approvals

All authors have read and approved the final version of the manuscript.

## Competing Interests

The authors declare no conflicts of interest

## Acknowledgments

The study was conducted at the unique research facility “Center for Neurocognitive Research (MEG-Center)” of MSUPE.

## Funding statement

This research was supported by the Russian Science Foundation (project # 25-18-00739).

## Notes

### Competing Interest Statement

The authors have declared no competing interest.

