## Supplementary figure S1 for "Top-down attention modulates auditory sustained responses but not the neural processing advantage for vowels"

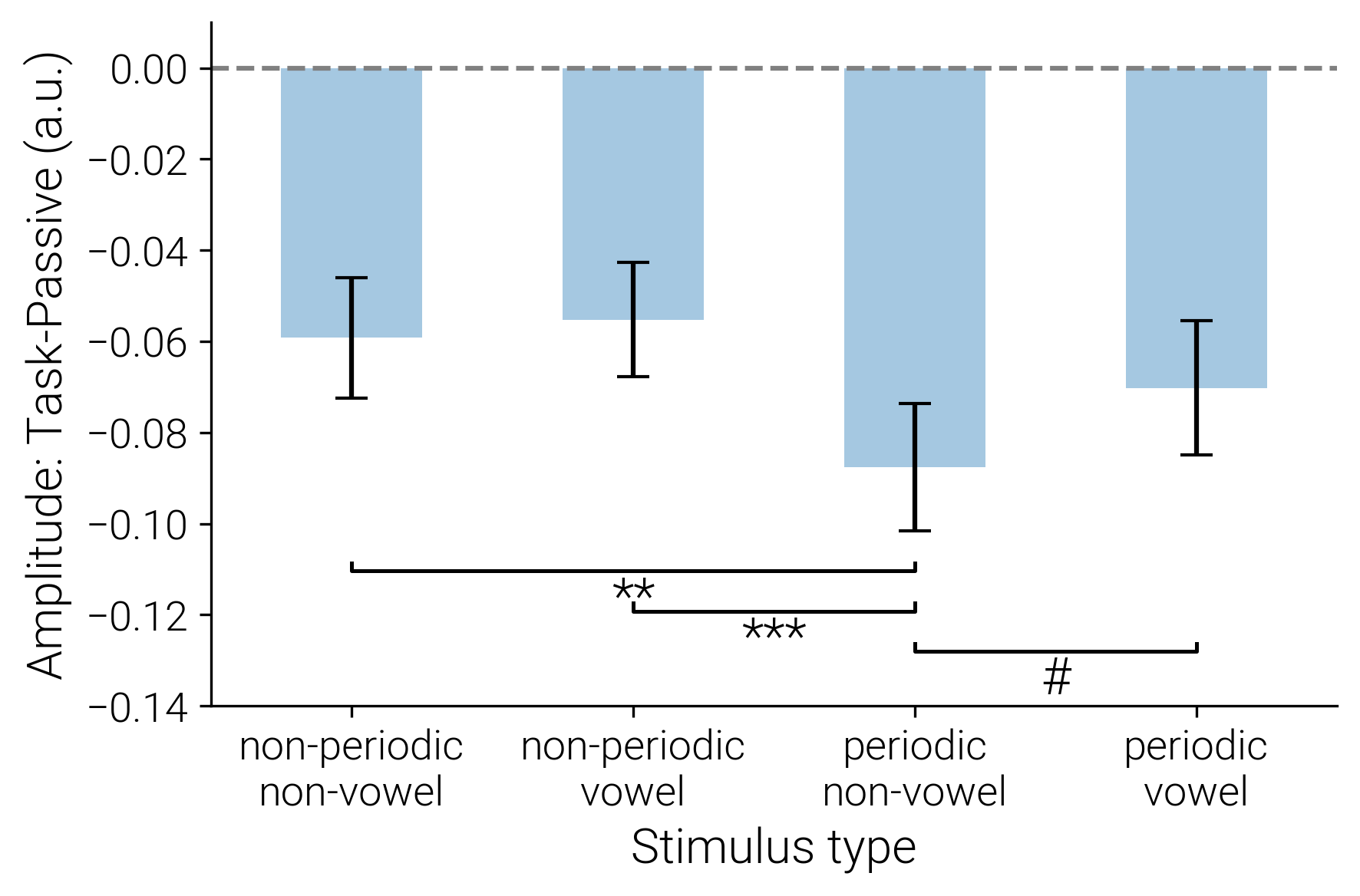

**Figure S1.** Change in the early portion (70-150 ms) Sustained Negativity (SN) during auditory task relative to passive listening (all Trial Positions combined). *** p<0.001; ** p<0.01; # p=0.07

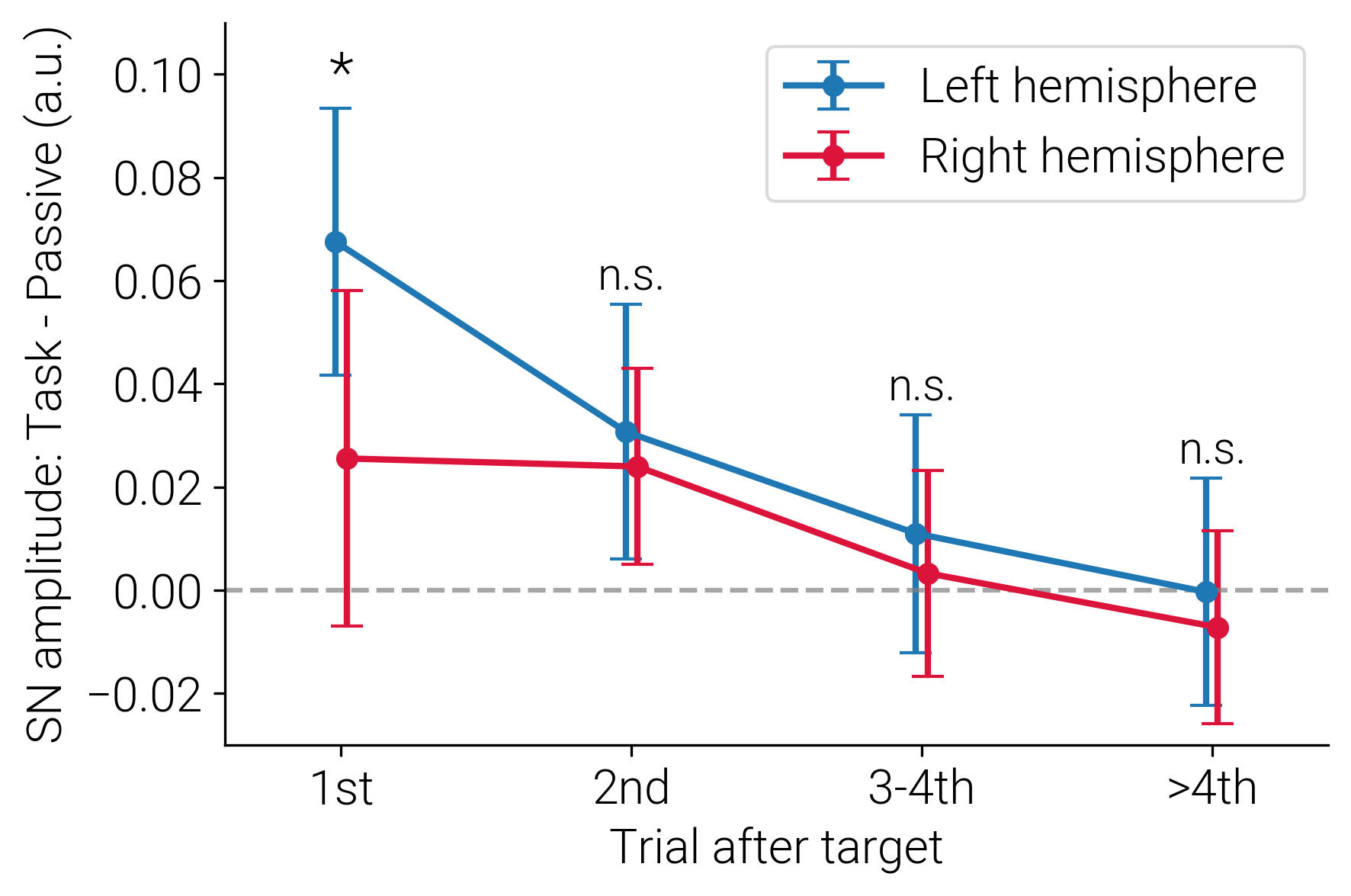

**Figure S2.** Change in the middle portion (150 - 250 ms) of Sustained Negativity (SN) in the left and right hemispheres during auditory task relative to passive listening (all Stimuli Types combined).
